# Comparative analyses of two primate species diverged by more than 60 million years show different rates but similar distribution of genome-wide UV repair events

**DOI:** 10.1101/2020.04.06.027201

**Authors:** Umit Akkose, Veysel Ogulcan Kaya, Laura Lindsey-Boltz, Zeynep Karagoz, Adam D. Brown, Peter A. Larsen, Anne D. Yoder, Aziz Sancar, Ogun Adebali

## Abstract

Nucleotide excision repair is the primary DNA repair mechanism that removes bulky DNA adducts such as UV-induced pyrimidine dimers. Correspondingly, genome-wide mapping of nucleotide excision repair with eXcision Repair sequencing (XR-seq), provides comprehensive profiling of DNA damage repair. A number of XR-seq experiments at a variety of conditions for different damage types revealed heterogenous repair in the human genome. Although human repair profiles were extensively studied, how repair maps vary between primates is yet to be investigated. Here, we characterized the genome-wide UV-induced damage repair in gray mouse lemur, *Microcebus murinus*, in comparison to human. Mouse lemurs are strictly nocturnal, are the world’s smallest living primates, and last shared a common ancestor with humans at least 60 million years ago. We derived fibroblast cell lines from mouse lemur, exposed them to UV irradiation. The following repair events were captured genome-wide through the XR-seq protocol. Mouse lemur repair profiles were analyzed in comparison to the equivalent human fibroblast datasets. We found that overall UV sensitivity, repair efficiency, and transcription-coupled repair levels differ between the two primates. Despite this, comparative analysis of human and mouse lemur fibroblasts revealed that genome-wide repair profiles of the homologous regions are highly correlated. This correlation is stronger for the highly expressed genes. With the inclusion of an additional XR-seq sample derived from another human cell line in the analysis, we found that fibroblasts of the two primates repair UV-induced DNA lesions in a more similar pattern than two distinct human cell lines do. Our results suggest that mouse lemurs and humans, and possibly primates in general, share a homologous repair mechanism as well as genomic variance distribution, albeit with their variable repair efficiency. This result also emphasizes the deep homologies of individual tissue types across the eukaryotic phylogeny.

## Introduction

Nucleotide excision repair is an essential mechanism to remove bulky DNA adducts including UV-induced DNA lesions (Hu et al. 2017b). As in other repair systems, excision repair starts with damage recognition. Two subpathways based upon damage recognition lead to two repair mechanisms: global repair (GR) and transcription-coupled repair (TCR). GR is active throughout the genome whereas TCR is only active on the transcribed strands as it is initiated by damage recognition through stalled RNA polymerase II (RNAPII). To date, many techniques have been developed to detect DNA damage and repair (Li and Sancar 2020). The approach of some methologies has been NGS-based, which allows answering genome-wide questions. To reveal genome-wide excision repair dynamics, heterogeneity and associations, eXcsion Repair sequencing (XR-seq) was developed. XR-seq technique directly measures the repair events by capturing the excised oligomer containing the lesion. It was found that TCR is more efficient particularly for slowly-repaired DNA lesions. For instance, among UV photoproducts 6-4 pyrimidine-pyrimidone ([6-4]PP) and cyclobutane pyrimidine dimer (CPD), CPD is more prone to TCR as it is less efficiently repaired by GR. Although DNA lesions might preferentially form at certain local sites in the genome, the overall heterogeneity of repair is majorly due to uneven repair efficiency throughout the genome (Hu et al. 2017a; Mao et al. 2018). Genomewide heterogenous repair distribution is mostly caused by transcription and chromatin structure (Hu et al. 2015, 2016; Adar et al. 2016; Yimit et al. 2019).

To date, genome-wide repair maps were generated for model organisms including *Escherichia coli* (Adebali et al. 2017a, 2017b), *Saccharomyces cerevisia* (Li et al. 2018), *Drosophila melanogaster* (Deger et al. 2019), *Arabidopsis thaliana* (Oztas et al. 2018), *Mus musculus* (Yang et al. 2018), and *Homo sapiens* (Hu et al. 2015, 2017a). TCR presence was verified for each of these species. For eukaryotic genomes, the consistent finding was the efficient repair in open chromatin regions. Heterochromatin regions were found to be repaired at later time points. Human repair profiles were extensively studied with respect to damage formation and chromatin states. Whether regions in human genome that are efficient with respect to repair are organism-specific is yet to be investigated.

To study whether repair patterns are unique to the organism of interest we aimed to compare human and a deeply diverged non-human primate. Gray mouse lemur (*Microcebus murinus*) stands out as a promising model organism candidate because of its small body size, short gestation time (2 months) and fast sexual maturation (6-8 months) (Blanco et al. 2015; Ezran et al. 2017). A near chromosome level reference genome for the gray mouse lemur was recently sequenced and assembled (Larsen et al. 2017). With no surprise, it was shown that mouse lemur and human orthologs share ~91% identity. Although a robust genome assembly is available, we lack an in-depth understanding of this species’ genomic features such as epigenetic maps, transcriptomes and methylomes.

Given that mouse lemurs and humans last shared a common ancestor at the base of the primate clade (Horvath and Willard 2007), same cell types from human and mouse lemur should behave similary in response to DNA damage as a reflection of their deep homology. With this motivation, we carried out a comparative study between these two primates to understand similarities and differences between their repair profiles. We derived primary fibroblasts from mouse lemur and immortalized them. We performed survival assays in response to UV stress, immunoslot blot assays and in vivo excision assays for both cell lines. From mouse lemur fibroblasts, we obtained transcriptomes, exposed cells to UV and performed XR-seq. XR-seq captured the excised oligomers as repair products for two main UV-induced damage types: (6-4)PP and CPD. We compared lemur and human fibroblast XR-seq datasets with respect to their genomic repair distribution.

## Results

### Excised oligomer characteristics

In vivo excision assay resulted in the excised oligomers containing UV-induced lesions (Fig 1A). Excised oligomers were captured for two distinct damage types with specific anti-damage antibodies. The oligomer length distribution varies from 16 to 30nt. Two intense bands were observed, which indicate primary and degraded excised oligomers. The gel images show that the primary excised oligomer lengths vary between 23-25nt. Through time course, gel images reflect a more intense secondary product band, suggesting higher levels of degraded excised oligomer with damage is observed at later time points. For normal human fibroblasts 1 (NHF1) and mouse lemur fibroblasts, this trend is similar.

**Figure 1:**
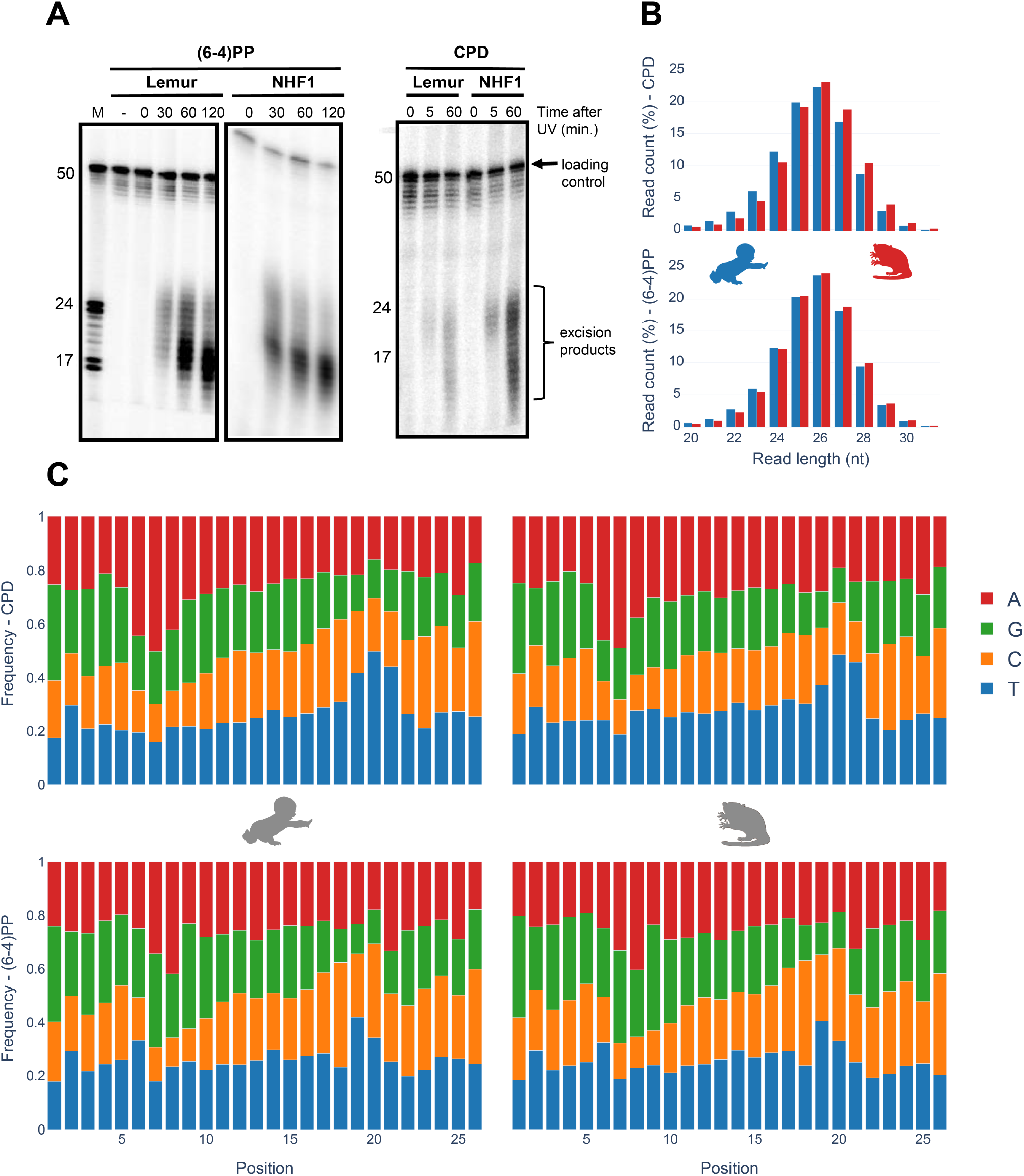
Characteristics of excised oligomers upon repair of UV-induced damages. *(A)* In vivo excision assay for (6-4)PP and CPD are shown. *(B)* XR-seq read length distribution of excised oligomers for CPD (top) and (6-4)PP (bottom). Human and mouse lemur are shown in blue and red, respectively. *(C)* Nucleotide frequency in the predominant oligomer (26nt) for each primate and damage type. Representative data from replicate 1.

Nucleotide excision repair in human, *S. cerevisiae* and *E. coli* was shown to yield primary excised oligomers predominantly in lengths of 27, 24 and 13 nucleotides, respectively (Hu et al. 2013; Huang et al. 1992; Li et al. 2018; Sancar and Rupp 1983; Adebali et al. 2017a). Mouse lemur XR-seq resulted in oligomer length distribution identical to humans (Fig 1B) thus indicating deep homology with human. For both CPD and (6-4)PP, nucleotide excision repair machinery incised 26nt oligomer as the predominant product. A small variance in oligomer lengths between in vivo excision assay and XR-seq might be due to the immediate exonuclease activity on excised oligomer for in vivo excision assay. Because of the TFIIH coimmunoprecipitation in excision repair sequencing technique, XR-seq primary products are not yet degraded, possibly because of TFIIH protection of the excised oligomers.

### Identical CPD and (6-4)PP nucleotide frequency between human and lemur

Nucleotide frequencies of the excised oligomers between human and mouse lemur are identical for both CPD and (6-4)PP (Fig 1C). Pyrimidine enrichment was revealed at 19th, 20th and 21st nucleotides for CPD and (6-4)PP for both organisms, which is in agreement with the incision site at 6 to 8nt fixed distance to the 3’ end of the DNA lesion. The nucleotide contents for CPD and (6-4)PP are different, as they were previously reported and discussed (Hu et al. 2015).

### UV repair efficiency of mouse lemur in comparison with human

To identify the overal repair efficieny upon UV irradiation we performed a colony survival assay. We seeded 300 cells and grew them for 16 hours. Cells were exposed to UV-C at 1 J/m^2^sec for variable time intervals to reach indicated doses ranging from 2.5 to 10 J/m^2^. We counted cells for each dose and plotted survival curves for lemur and human fibroblasts (Fig 2A). Based on the survival curves we infer that mouse lemur fibroblasts are more sensitive to UV relative to human cells. This suggests a differential repair rate between the two species which is possibly reflective of the diurnal versus nocturnal habits of humans and mouse lemurs, respectively.

**Figure 2:**
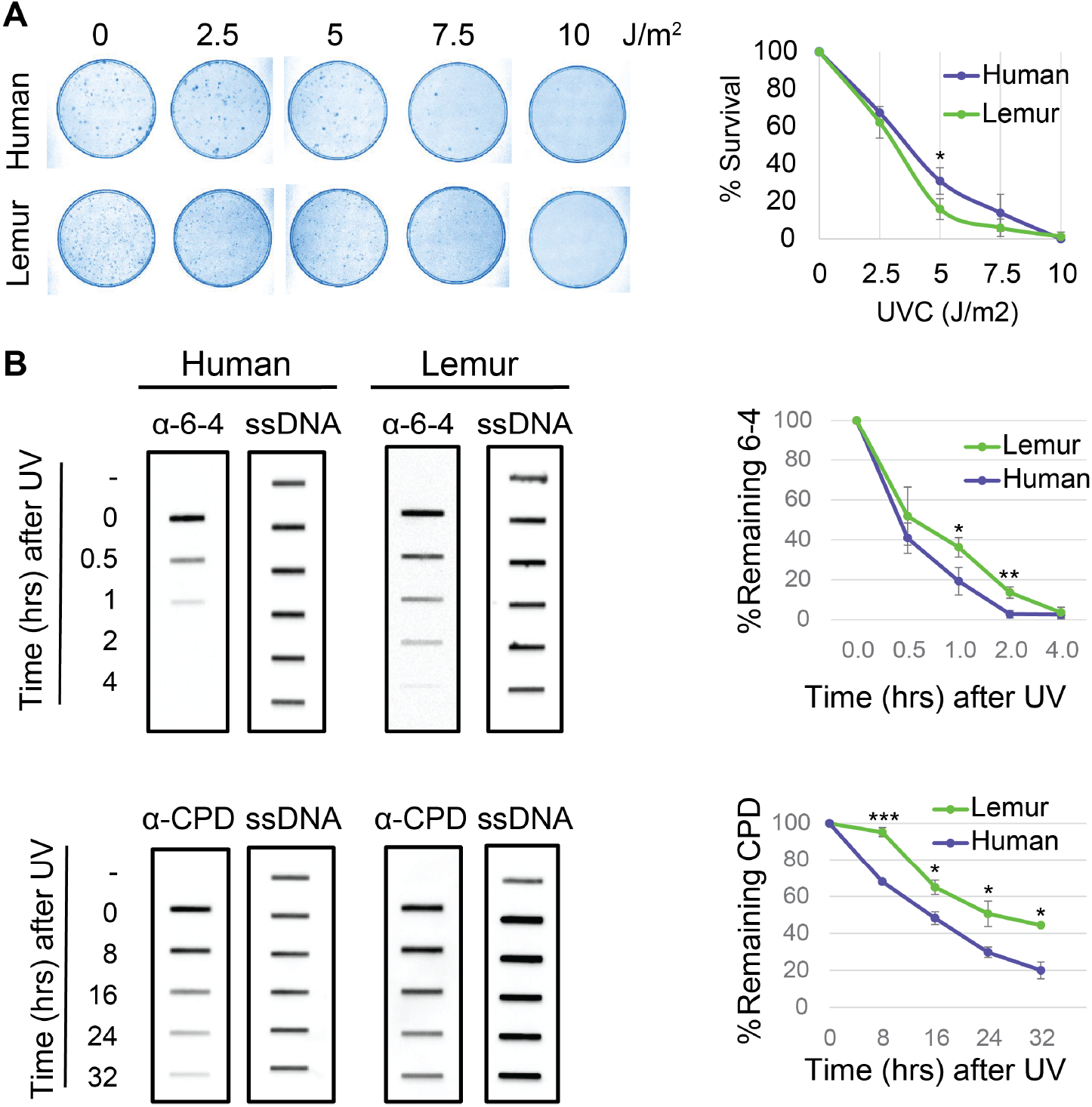
UV sensitivity and repair rates in human and lemur fibroblasts. *(A)* Clonogenic survival assay (left) and the quantified results (right). *(B)* Immunoslot blot repair assays to examine the repair kinetics of UV-induced DNA lesions for both organisms and each damage type (left). Remaining damage levels at each time point were quantified and plotted (right). All experiments were repeated three times, and graphed data are presented as (mean+/-SD). Tests (t-test) performed (H_0_:μ_human_=μ_lemur_) for each dosage (A) and time points (B). * P ≤ 0.05; ** P ≤ 0.01; *** P ≤ 0.001; rest P > 0.05.

We then performed immunoslot blot assays (Fig 2B). Cells were exposed to UV-C at 10J/m^2^. To separate UV-induced DNA lesions we used specific antibodies for CPD and (6-4)PP. We quantified the remaining damage signal for each damage type for both species. Both (6-4)PP and CPD repair appeared to be more efficient in human cells compared to mouse lemur. Human (6-4)PP repair was completed in 2 hours whereas in lemurs, it took 4 hours to complete (6-4)PP repair. 20% of CPDs were still unrepaired for human cells 32 hours after UV irradiation. Unrepaired CPD ratio was more than 40% for mouse lemur cells. Survival and immonoslot blot assays are in harmony and they suggest mouse lemur cells have relatively low repair rates compared to NHF1 cells.

### Prominent transcription-coupled repair in mouse lemur

Nucleotide excision repair comprises two sub-pathways each of which distinctly acts on damage recognition. GR is the genome-wide mechanism that is active in any region in the genome, although its efficiency depends on the type of damage as well as chromatin factors. On the other hand, TCR depends on the RNAPII stalled at the lesion. Stalled RNAPII recruits transcription-coupled repair factor CSB to enhance repair. As TCR occurs only when RNAPII is stalled, it is solely active in the regions that are transcribed by this polymerase. Also, TCR is seen in the transcribed strand of the genes. DNA lesions in non-transcribed strands are subject to global repair. A major rate-limiting step of nucleotide excision repair is damage recognition. Therefore, minor helix-distorting lesions such as CPDs are recognized more efficiently when they stall otherwise elongating RNAPII. In those organisms having TCR, genes have asymmetrical repair between two strands; transcribed strand (TS) is repaired more efficiently compared to nontranscribed strand (NTS) (Hanawalt and Spivak 2008; Mellon and Hanawalt 1989).

Global repair for (6-4)PP is much faster than for CPD (Steurer et al. 2019; Adar et al. 2016). TCR removes a minor fraction of the (6-4)PPs, and thus strand asymmetry between TS and NTS (TS/NTS ratio) is much weaker for (6-4)PP compared to TS-favored strand asymmetry of CPD repair (Adar et al. 2016). CPDs, on the other hand, are harder to be recognized by global repair, and therefore it takes a longer time to remove CPDs from the genome. For this reason, at early time points (such as 1 hour after UV irradiation), we observe a strong TCR effect which is indicated by the asymmetrical repair between TS and NTS. Here, we compared the genic strand asymmetry between human and mouse lemur (Fig 3). We retrieved all annotated protein-coding genes from both genomes. We removed the genes that are closer to each other with less than 20kb in order to remove the signal that might have a “canceling-out” effect. We used 5277 and 3366 annotated genes for human and mouse lemur, respectively (see methods). We performed a meta-analysis where we aligned all transcription start and transcription end sites and calculated the RPKM (reads per kilobase per million mapped reads) values. Interestingly, mouse lemur fibroblasts exhibited stronger TCR profiles compared to human cell lines at 1 hour after CPD formation (TS/NTS Mann-Whitney U test p=4.3*10^−21^).

**Figure 3:**
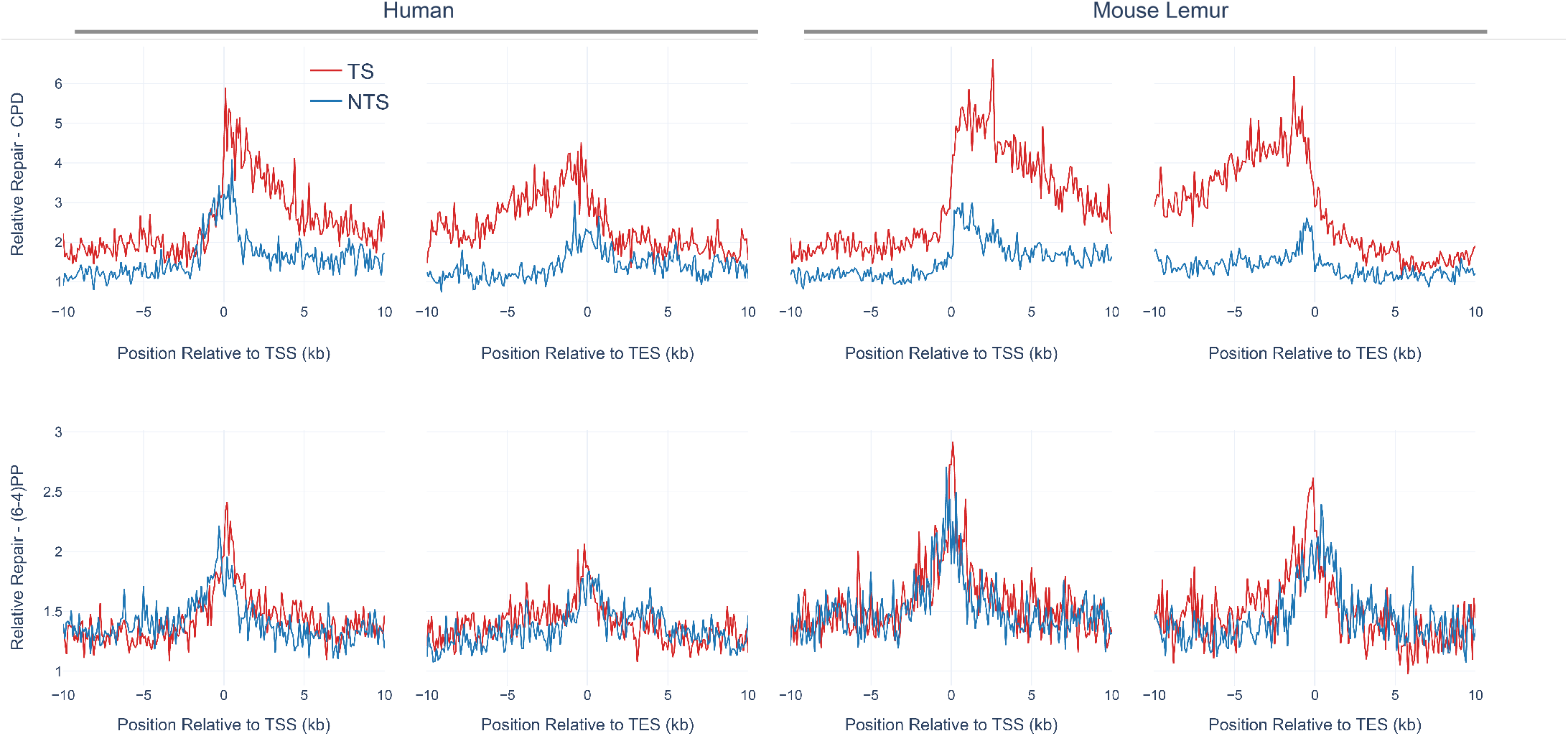
Transcription-coupled repair in mouse lemur and human fibroblasts. Average profiles of CPD XR-seq (top) and (6-4)PP XR-seq (bottom) over 5277 and 3366 annotated genes are plotted for human (left) and mouse lemur (right), respectively. Transcription start sites (TSS) and transcription end sites (TES) were retrieved from GTF (Gene Transfer Format) files for the two genomes. Transcribed (red) and nontranscribed (blue) strand repair are shown in the downstream and upstream of TSS and TES, respectively. 10 kb upstream and downstream of TSS and TES were divided into 100 bp windows. XR-seq reads aligned to each bin were normalized to RPKM (reads per kilobase per million mapped reads). The XR-seq RPKM values were normalized by the shuffled RPKM values derived by mapped reads aligned to random genomic sites. Only replicate 1 is shown. TS/NTS median values for TSS downstream for CPD are 1.76 and 2.31 for human and mouse lemur, respectively.

### Conserved repair rates between homologous regions of two primates

In order to compare and contrast the genome-wide repair profiles between human and mouse lemur, we retrieved the homologous genomic regions between the organisms (Fig 4A). We aligned the two genomes and kept the regions that have at least 80% identity. Due to the low coverage of the XR-seq samples we removed the aligned segments that are shorter than 400 bases to eliminate the random effect due to scarce repair events. Lineage-specific duplications (or deletions) result in paralogs that can be co-orthologs to one locus in the other genome. This situation is known as the “one-to-multiple” orthology relationship. As we cannot be sure which one of the multiple paralogs in one genome is the “true” ortholog of a region in the other one, we filtered out one-to-multiple kinds of homologous relationships and kept one-to-one homologs only.

**Figure 4:**
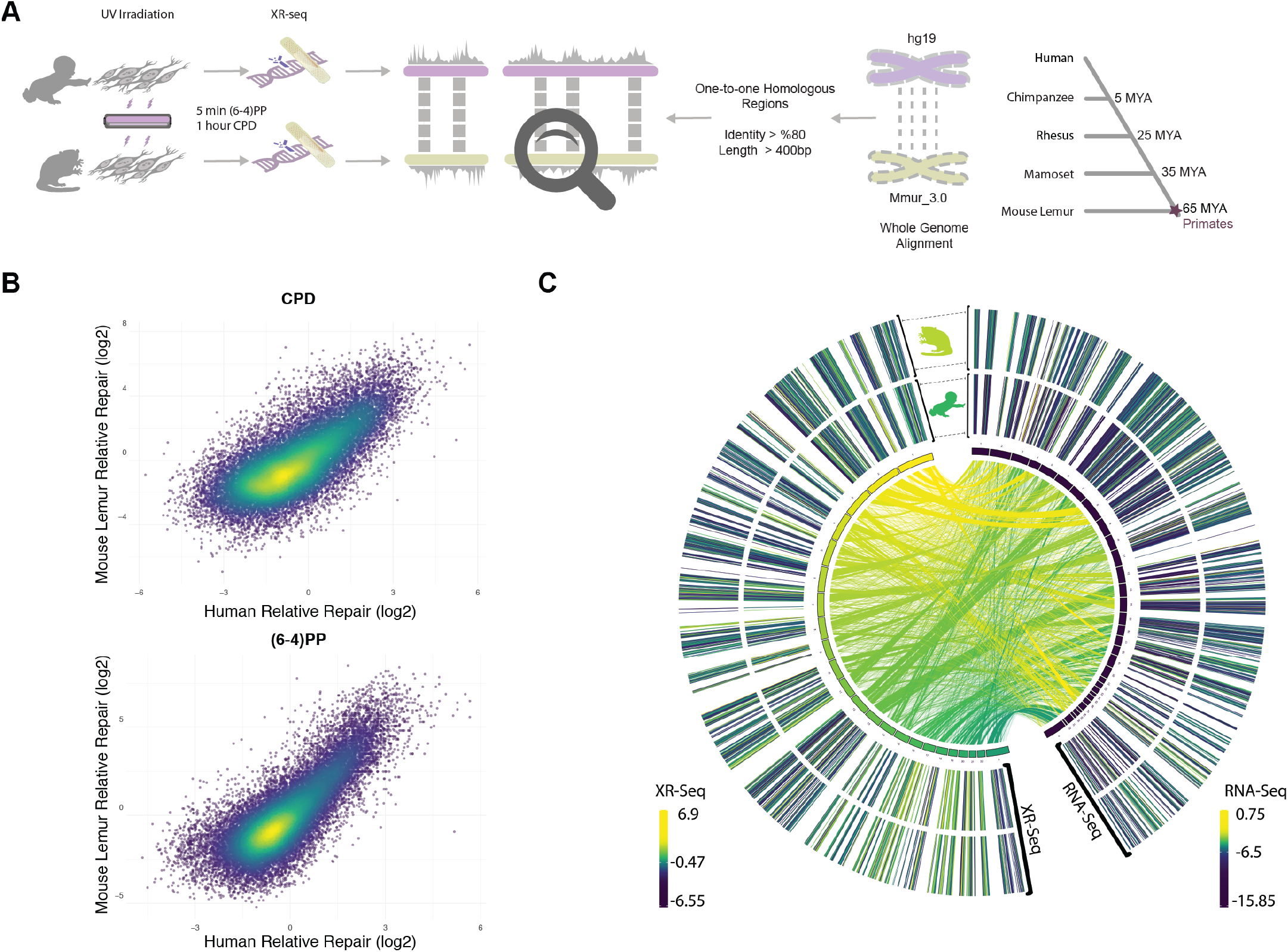
Comparative analysis methodology and correlated repair rates between human and mouse lemur. A) Approach to identify orthologous regions between human and mouse lemur (see methods for details). B) Scatter plots showing the normalized repair levels between two organisms. Relative repair was calculated by normalizing the repair signal by the simulated XR-seq reads to eliminate the sequence context bias. C) Mapped orthologous regions with repair and transcription profiles mapped at left and right outer rings, respectively. Inner circle represents the human (left) and mouse lemur (right) chromosomes. Inner lines connecting orthologous regions are colored based human chromosome color scale. Repair and transcription values (outer rings) for genomic regions that have no evident ortholog (based on the criteria in A) are not shown.

1.0% and 1.3% of the genomes were taken into account with one-to-one homologs for human and mouse lemur, respectively. 4-5% and 2-3% of the XR-seq reads mapped to those regions for CPD and (6-4)PP, respectively. The reason why the mapping rate of CPD is higher than (6-4)PP could be due to its slower repair rate, as slowly repaired CPDs are more prone to the effects of chromatin. Because we applied stringent criteria to retrieve the homologous regions with at least 80% identity, they are likely to be more open compared to other regions in the genome. Therefore, more CPD repair events aligned to those regions relative to (6-4)PP reads.

We investigated the correlation between the repair profiles of the two species. Because excised oligomers have a certain nucleotide bias due to the damage site, we normalized the true repair signal (XR-seq) by the simulated XR-seq reads. Simulated reads have the nucleotide content of the true XR-seq reads (Fig S1), however they are randomly retrieved from the homologous regions. With this approach, we eliminated potential bias of the nucleotide content that creates a pseudo-correlation. Although there is a clear difference in the relative TCR rates between two species (Fig 3), homologous regions of two genomes exhibited a strong repair rate correlation; R values for (6-4)PP and CPD are 0.80 and 0.72 (p=0), respectively (Fig 4B). Not only repair, but also transcription levels between the two species are correlated (Fig 4C). The similar repair and transcription profiles between two cell lines are independent of their chromosomal locations. Chromosomal homologous region associations are dispersed as previously suggested (Larsen et al. 2017).

### Repair profile correlation is associated with gene expression

To test whether gene expression is a factor associated with the correlation of repair between these two highly-diverged species of primates, we examined the correlation strength and gene transcription levels. We prepared RNA-seq data sets for mouse lemur fibroblasts and obtained RNA-seq datasets for human fibroblasts from a publicly available database (see methods). The strength of the correlation between the two repair profiles was found to be correlated with transcription levels (Fig 5). Highly expressed regions had better repair correlation between the two species (Figs S2;S3;S4;S5).

**Figure 5:**
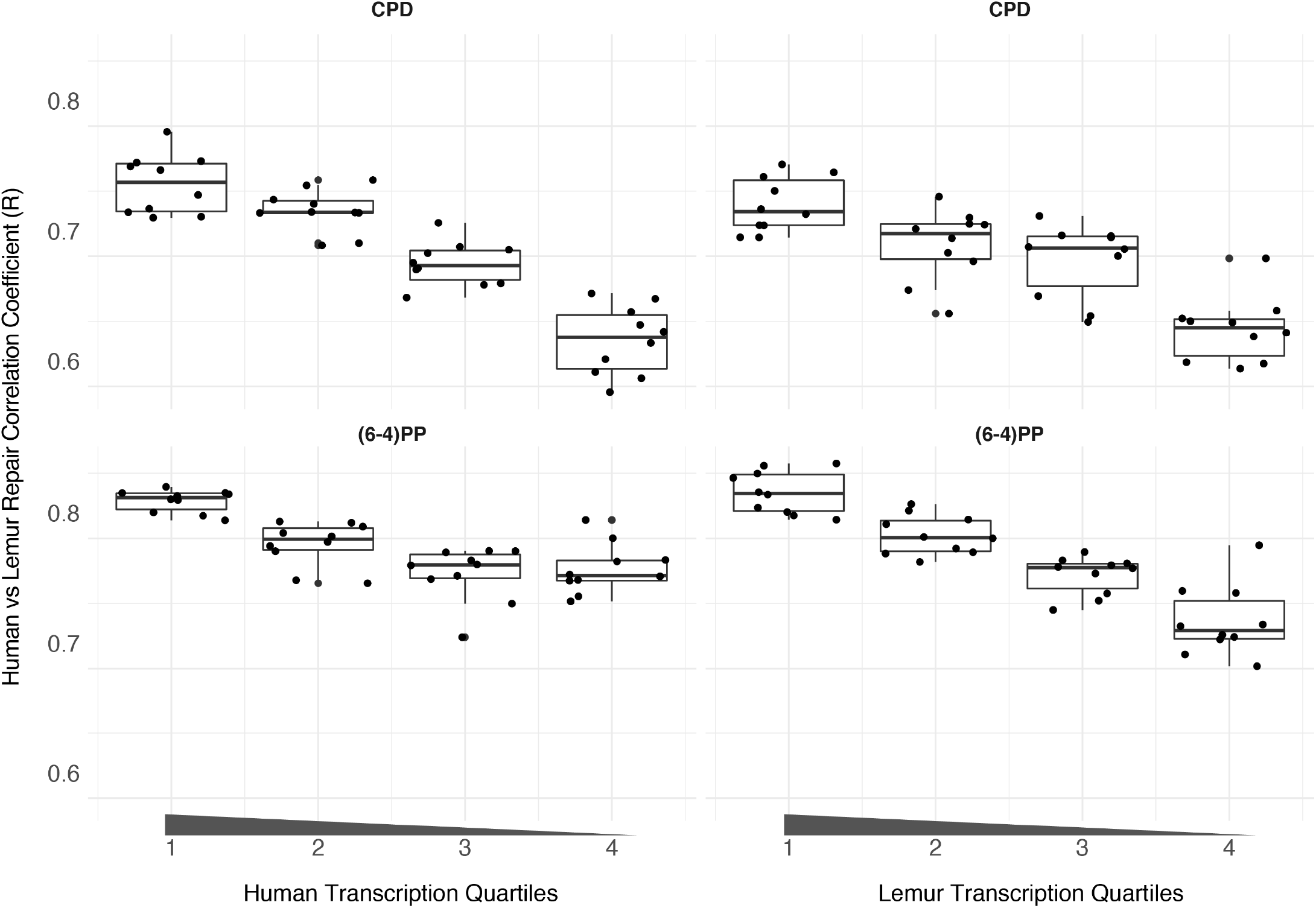
Repair rate consistency in correlation with gene expression. The transcription levels of the orthologous regions were divided into quartiles separately for human and lemur. Out of the quartiles, 10 bootstrapped subsamples were retrieved, and for those regions repair correlation was analyzed. Correlation coefficient (R values) are represented on the y-axis; quartiles are on the x-axis.

### The same cell type from two primates have more similar repair profiles relative to two different cell types from the same organism

We cannot dismiss the possibility that the correlation of repair profiles between two primates could simply be due to the heterogenous mappability of the genomes. Such a correlation makes sense only when we measure the similarity of repair patterns relative to another dataset. For this reason, we used another human cell type, GM12878 - B-Lymphocytes to ask whether the repair profile of human fibroblasts is closer to lemur fibroblasts or to human lymphocytes. We performed XR-seq for CPD as it yields relatively high genome-wide heterogeneity compared to (6-4)PP. Although repair profiles between human GM12878 and NHF1 are also correlated (Fig 6A), this correlation is not as strong as the one between human and mouse lemur fibroblasts. Additionally, human and mouse lemur fibroblasts clustered together in the PCA plot (Fig 6B). Within fibroblasts, biological replicates grouped together. More interestingly, (6-4)PP and CPD samples formed distinct groups each of which has human and mouse lemur samples in the subclusters. The two replicates of CPD XR-seq samples of human lymphocytes (GM12878) clustered as an outgroup. This result suggests that human and mouse lemur fibroblasts have similar chromatin and expression patterns and therefore have similar repair profiles. On the other hand, even though derived from the same organism, completely different human cell lines have a considerable variation with respect to repair patterns. It is also interesting to note that although TCR profiles between two primates are quite different from each other, they share high similarity in repair profiles.

**Figure 6:**
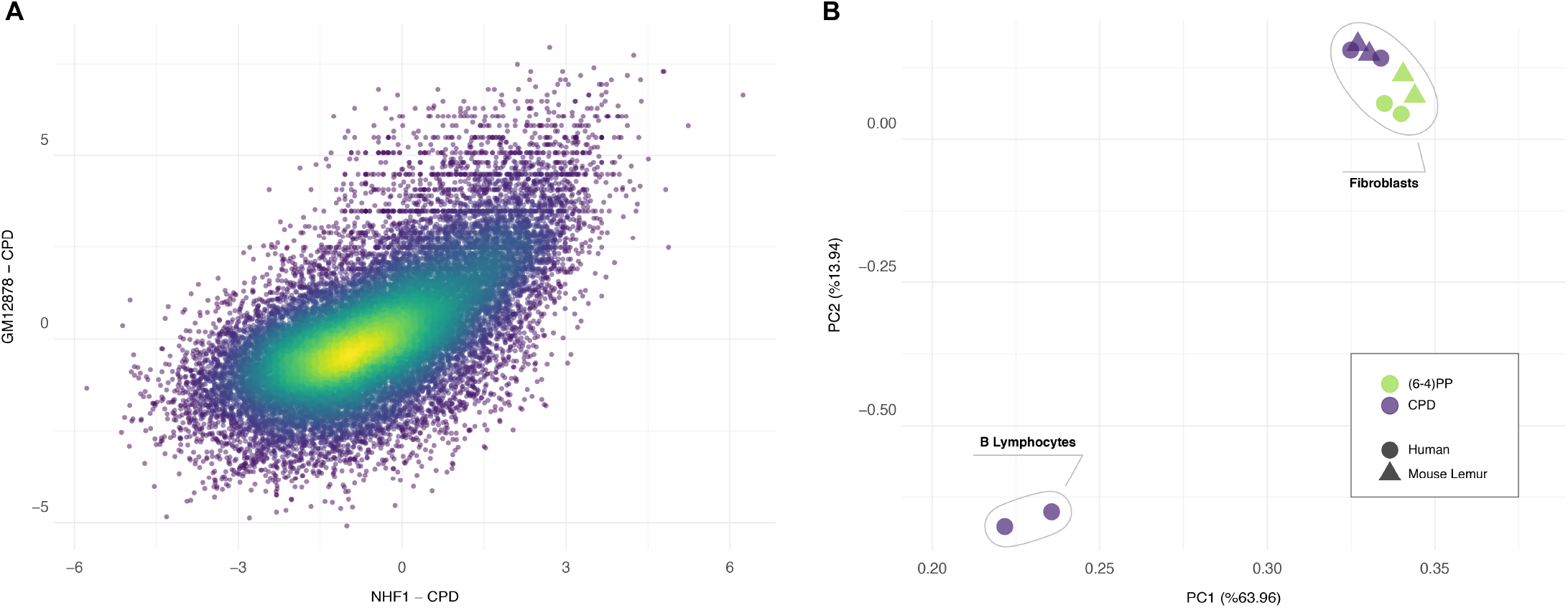
Cell type-based consistency of repair profiles between two primates. A) Scatter plot or normalized repair levels between two human cell lines, GM18787 and NHF1, show correlation (R=0.65, p=0). B) Principle component analysis of 10 XR-seq samples.

## Discussion

XR-seq methodology has become prominent in genome-wide profiling of nucleotide excision repair at single nucleotide resolution. This technique has been applied to a variety of model organisms from human to bacteria. Although genome-level understanding of human repair behavior is relatively well established in the recent years, the associations between closely related primates with respect to UV damage response and genome-wide repair distribution were entirely lacking. With this study, we established the differences and similarities between mouse lemur and human. First, we observed repair efficiency differences between the two genomes of the two major types of DNA damage induced by UV. Like in humans, (6-4)PP was much more quickly completed compared to CPD in mouse lemur. Both (6-4)PP and CPD repair efficiencies were apparently slower in mouse lemur fibroblasts relative to NHF1 cells. This observation is in agreement with the survival assays which show that mouse lemur cell lines were more sensitive to UV, which contains both damage types we studied. And although this comparison is limited to only two species at present, we postulate that these differences could relate to the diurnal and nocturnal lifestyles of the two species. Given that mouse lemurs are rarely exposed to UV light in nature, there may be some physiological relevance that their UV-induced damage repair would be less efficient than that seen in humans.

Interestingly, we also observed that TCR was relatively more efficient in mouse lemurs compared to humans especially for CPD repair. TCR of (6-4)PP was not prominent, as was previously shown, and our immunoslot blot assays indicate that (6-4)PP repair is fast and therefore mostly repaired by GR rather than TCR. We observed similar TS and NTS repair signal for (6-4)PP in genic regions, also suggesting that GR is the primary subpathway to repair (6-4)PP. A striking fact is the inverse correlation between TCR, which is measured by TS/NTS repair signal, and overall repair efficiencies. Mouse lemurs with more efficient TCR falls behind in overall repair compared to human. A possible explanation for this observation is that global repair is the main determinant of the repair efficiency. XR-seq yields the entire nucleotide excision repair events with no distinction between TCR and GR. For this reason, GR and TCR efficiencies derived thourgh XR-seq are relative to each other. Like the relativity in any RNA-seq, which cannot measure the total transcription levels, XR-seq also gives the relative distribution of the repair events. Therefore, measured “TCR efficiency” actually corresponds to “GR deficiency” in XR-seq profile. In other words, when the damage is not efficiently recognized by GR pathway, TCR takes over the recognition and removes DNA lesions in TS. With this study, previously observed high TCR levels in *Arabidopsis thaliana* (Oztas et al. 2018) and mouse (Yang et al. 2018) now also suggest relatively poor GR efficiency in those organisms.

Human evolution has been under investigation since the time of Charles Darwin. Comparative analyses between human and other animals have revealed both shared and human-specific features. In the era of molecular biology it has been possible to reveal molecular features as they are shared between humans and their relatives. Primates are an extremely diverse group comprised of species that range in body size from 30-80 grams (mouse lemurs) to 150kg (African apes). Mouse lemurs stand out as the world’s smallest living primate. Given detailed knowledge of primate phylogeny and divergence times (Reis et al. 2018), we know that the most recent common ancestor (MRCA) of humans and mouse lemurs is MRCA of all living primate species, and that this MRCA likely arose near the Cretaceous-Paleogene (K-Pg) boundary 65 mya. Thus, we can reasonably conclude that features that are shared between humans and mouse lemurs are homologous and characteristic of the primate clade.

We have a limited knowledge on the epigenetic architecture of mouse lemurs as we far lack chromatin data. Prior to this study, it was unknown whether repair profiles would be similar in the comparison of two primates species. Our work revealed that human and mouse lemur fibroblasts are significantly similar with respect to their preference on which genomic region to repair first. This bias is likely to be caused by the chromatin structure as previously shown in other organisms (Yimit et al. 2019). Damage distribution is known to be homogenous at large genomic scales (Hu et al. 2017a). Therefore, it is safe to assume that repair preference is the major factor for XR-seq heterogeneity. The repair profiles of human and mouse lemur fibroblasts are much more similar to each other than the ones between two distinct human cell lines. Moreover, (6-4)PP and CPD repair profiles are more similar to themselves, suggesting similar repair dynamics between two organisms. Further studies on mouse lemur epigenetics will possibly explain the most differentiated genomic repair levels in mouse lemurs. Better understanding mouse lemur genomics, as well as other species within the strepsirrhine clade, will provide additional insight into human evolution, repair and mutagenesis.

## Methods

### Cell lines and reagents

#### Establishment of the immortalized mouse lemur fibroblast cell line

Primary fibroblasts were established using methods outlined in Larsen and Harris et al (Larsen et al. 2017). For the current study, we used an immortalized version of these primary mouse lemur fibroblasts. This cell line was generated by transfecting passage six primary cells at approximately 60% confluency with 5 μg of human hTERT (hTERT) plasmid DNA per well of a 6-well plate using Lipofectamine^®^ 2000 diluted in Opti-MEM^®^ medium. Twenty-four hours post transfection, cells were placed under selection using G418 sulfate at a concentration of 300 μg/mL for two weeks. Selected cells were then plated into a 96-well plate and remained under G418 sulfate maintenance at a concentration of 100 μg/mL to establish single cell colonies. Approximately 2 weeks post transfection, colonies were observed in 19 of the 96 wells and these cells underwent subsequent propagation and expansion.

#### Normal human fibroblasts

Telomerase-immortalized normal human fibroblasts (NHF1) were previously described (Heffernan et al. 2002; Hu et al. 2015). All cells were grown in Dulbecco’s modified Eagle’s medium supplemented with 10% fetal bovine serum at 37°C in a 5% CO2 humidified chamber.

### Clonogenic Survival Assay

Cells were seeded in 10 cm plates at 300 cells/plate, grown for 16 hours, and then UV irradiated as follows. After removing and reserving culture medium, cells were placed under a GE germicidal lamp that emits primarily 254-nm UV light (UV-C) at 1 J/m^2^sec connected to a digital timer to give the indicated dose. Following irradiation, the culture medium was replaced, and the cells were incubated for 7 days. Plates were then washed with phosphate-buffered saline (PBS), the cells were fixed for 10 min in 75% methanol/25% acetic acid, and stained with 0.5% crystal violet stain (J.T. Baker) in 25% methanol for 30 min. The plates were then washed extensively in tap water to remove the excess stain. After drying, images of the stained cells were taken on a Molecular Imager Chemi-Doc XRS+ system (Bio-Rad) for presentation of representative, qualitative results. Colonies were counted and the surviving fraction was calculated by dividing the number of colonies on treated plates by the number on the untreated plates. Each condition was performed in triplicate and the experiment was repeated three independent times.

### Immunoslot Blot Analysis

Repair of UV photoproducts from genomic DNA was measured as described previously (Gaddameedhi et al. 2010). Cells were UV irradiated as described above for a dose of 10J/m^2^. Following irradiation, the culture medium was replaced, and the cells were incubated for the indicated periods of time. Genomic DNA (250 ng), isolated with a QIAamp DNA Mini kit (Qiagen), was immobilized on a nitrocellulose membrane with a Bio-Dot SF Cell immunoslot blot apparatus (Bio-Rad) and heated for 90 min at 80 °C under vacuum. Blots were blocked in 5% milk and probed with an anti-(6-4)PP antibody (Cosmo Bio 64M-2 cat#NM-DND-001) or anti-CPD antibody (Cosmo Bio TDM-2 cat#NM-DND-002) as indicated. The secondary antibody was horseradish peroxidase-linked anti-mouse IgG from GE Healthcare (catalog no. NA931V) and the chemiluminescent signals were visualized with Clarity Western ECL Substrate (Bio-Rad) and using a Molecular Imager Chemi-Doc XRS+ system (Bio-Rad). Following immunoblotting, the blots were re-blotted with anti-ssDNA antibody (Millipore MAB3034 clone 16-19) to ensure equal loading of DNA. The experiment was repeated three times, and representative results are presented.

### Detection of Excised Oligonucleotide Products of Nucleotide Excision Repair

Nucleotide excision repair activity was visualized as previously described (Hu et al. 2013). Cells grown to ~80% confluency in 15-cm plates were harvested the indicated time after irradiation with 20 J/m^2^ of UV. The cells were pelleted by centrifugation, subjected to a modified Hirt procedure where cell pellets were resuspended in a 10× packed cell volume of lysis buffer (50 mM Tris-Cl, pH 8.0, 10 mM EDTA, 1.2% SDS and 100 μg/ml RNase A) and incubated for 15 min at room temperature. Following addition of a one-fourth volume of 5 M NaCl, the mixtures were gently mixed and incubated on ice for >16 h. After centrifugation at maximum speed (20,000 × g) for 1 h, the supernatants were gently transferred to new tubes and treated with 20 μg of proteinase K for 15 min at 55 °C, extracted with phenol/chloroform, and then precipitated with ethanol. The pellet was washed with 500 μl of 70% ethanol and resuspended in 10 μl of buffer EB (10 mM Tris-Cl (pH8.5)). The excised oligonucleotide products of nucleotide excision repair were purified from the remaining material with either anti-(6-4)PP or anti-CPD antibodies as follows: For each reaction, 5 μl of protein G Dynabeads (Invitrogen, catalog no. 10003D) slurry and 5 μl of anti-rabbit Dynabeads (Invitrogen, catalog no. 11203D) slurry were washed three times with 50 μl of wash buffer I (20 mM Tris-Cl (pH 8.0), 2 mM EDTA, 150 mM NaCl, 1% Triton X-100, and 0.1% SDS) and then incubated with 1 μl of rabbit anti-mouse IgG and 1 μl of anti-(6-4)PP or anti-CPD antibody in 20 μl of IP buffer (20 mM Tris-Cl (pH 8.0), 2 mM EDTA, 150 mM NaCl, 1% Triton X-100, and 0.5% sodium deoxycholate) for 3 h at 4 °C. After incubation, beads were separated from the liquid with a magnet and then mixed with 100 μl of IP buffer and 10 μl of DNA. The mixtures were rotated at 4 °C overnight. The beads were then washed sequentially with 200 μl each of wash buffer I, wash buffer II (20 mM Tris-Cl (pH 8.0), 2 mM EDTA, 500 mM NaCl, 1% Triton X-100, and 0.1% SDS), wash buffer III (10 mM Tris-Cl (pH 8.0), 1 mM EDTA, 150 mM LiCl, 1% Nonidet P-40, and 1% sodium deoxycholate), wash buffer IV (100 mM Tris-Cl (pH 8.0), 1 mM EDTA, 500 mM LiCl, 1% Nonidet P-40, and 1% sodium deoxycholate) and finally twice with TE (10 mM Tris-Cl (pH 8.0) and 1 mM EDTA). The oligonucleotides containing UV photoproducts were eluted by incubation with 100 μl of elution buffer (50 mM NaHCO3, 1% SDS, and 20 μg/ml glycogen) at 65 °C for 15 min. The eluted DNA was then isolated by phenol/chloroform extraction and followed by ethanol precipitation. The excised oligonucleotides were resuspended in 10 μl of water, and half of the DNA was 3’-end labeled for 1 h at 37 °C in a 10-μl reaction containing 6 units of terminal deoxynucleotidyl transferase (New England Biolabs), 0.25 mM CoCl2, and 1 μCi of [α-32P]-3’-deoxyadenosine 5’-triphosphate (cordycepin 5’-triphosphate, Perkin Elmer Life Sciences) in 1× terminal deoxynucleotidyl transferase buffer (New England Biolabs). 2.5 fmol of a 50-nucleotide oligomer was included in all reactions as an internal control, and oligonucleotides of known length were resolved on all gels as size markers. Following phenolchloroform extraction and ethanol precipitation, the DNA was separated on urea-containing polyacrylamide gels, detected with a phosphorimager and was quantified using ImageQuant software (version 5.2, GE Healthcare). The experiment was repeated three times.

### XR-seq library preparation

Samples were processed for XR-seq as previously described (Hu et al. 2018). Cells grown to ~80% confluency in 15-cm plates were harvested either 5 minutes or 1 h after irradiation with 20 J/m^2^ of UV, depending on the damage, (6-4)PP or CPD respectively, to be analyzed. Samples from four plates were pooled, lysed, and immunoprecipitated with anti-TFIIH antibodies p62 (Santa Cruz Biotechnology, sc292) and p89 (XPB, Santa Cruz Biotechnology, sc293), and then ligated to adapters and processed for sequencing.

### RNA-seq library preparation

Total RNA was isolated from one 10-cm plate of exponentially growing NHF1 or Lemur cells using Trizol (Invitrogen) and the RNeasy Mini Kit (Qiagen) following the manufacturer’s instructions. The library preparation and strand-specific paired-end sequencing (2 × 150 bp) on a HiSeq 4000 platform (Illumina) was performed by Novogene Co., Ltd.

### Human and lemur genome alignment

MUMmer version 3.23 (Kurtz et al. 2004) was used to align grey mouse lemur genome (Mmur_3.0) to human genome (hg19) with `nucmer` subprogram. The alignments were filtered based on criteria taking length and identity into account. Alignments were listed in “delta format” by default by Nucmer. For every reference (human) - query (lemur) pair, we kept the alignments which form the longest mutually consistent set. Filtering step was performed with `delta-filter` subprogram. `show-coords` subprogram was used to display summary information such as position, percent identity and other features of each alignment, in Btab format with `-B` and `-rclo` arguements.

We used `generateBED.R` (Akkose et al. 2019) to write homologous human-lemur regions in a bed file. Due to low genome coverage in XR-seq data sets small regions introduce random repair values. Since we were not interested in too short alignments, we filtered out alignments shorter than 400 bp while generating the bed file. Additionally, alignments with at percent similarity lower than 80 were filtered out of the list. Resulting was a BED file with each homologus region between two genomes.

This output file contained regions in either genome that align to multiple regions in the other genome. To remove such regions, we used, we used `bedtools intersect` (Quinlan and Hall 2010) to intersect the file with itself. The command line we used was: `bedtools intersect -wo -s -a humanOverlapsLemur_short_noFilter.bed -b humanOverlapsLemur_short_noFilter. bed > human_intersect.txt`. We compared this intersected output file to the `bed` file generated previously using `findDupAln.R` (Akkose et al. 2019) to exclude regions in both genomes that align to multiple regions in other genome. This resulted in a `bed` file of one-to-one homologous regions without multiple alignments for one region.

### XR-seq analysis

We trimmed 3’ adapter sequences (TGGAATTCTCGGGTGCCAAGGAACTCCAGTNNNNNNACGATCTCGTATGCCGTCTTCTGCTTG) using Cutadapt (Martin 2011). Bowtie2 version 2.3.4.1 (Langmead and Salzberg 2012) was used to align sequencing reads onto reference genomes with default parameters. With a two-step conversion process, the `sam` format was first converted (with SAMtools (Li et al. 2009)) to `bam` format, which was then converted to `bed` format with BEDTools (Quinlan and Hall 2010). The resulting `bed` files were sorted by coordinates and duplicate regions were removed. Command line we used was: `sort -u -k1,1 -k2,2n -k3,3n ${SAMPLE}_cutadapt.bed >${SAMPLE}_cutadapt_sorted.bed`

### RNA-seq analysis

Sequencing reads were aligned to reference genome using STAR Aligner 2.6.1a (Dobin et al. 2012) with default parameters. The aligned reads were converted to `bed` format with `bedtools bamtobed`. The resulting `bed` files were sorted by coordinates and duplicate regions were removed. Command line we used was: `sort -u -k1,1 -k2,2n -k3,3n ${SAMPLE}.bed >${SAMPLE}_sorted.bed`

### XR-seq simulation

We evaluated the correlation of repair mechanisms with both simulated and real XR-seq datasets. Simulated datasets were generated for the overlapping regions between grey mouse lemur genome and human genome using ART simulator (Huang et al. 2011). We used `bedtools getfasta` program with the `bed` file containing homologus regions between two organisms to generate a reference `fasta` file for ART simulator to produce synthetic reads. By default ART simulator produces a `fastq` file with fiat nucleotide distribution. For a better representation of XR-seq characteristics in simulated dataset, we obtain nucleotide distribution frequencies of both species XR-seq reads. We applied a scoring function where each nucleotide in the simulated read was scored based on the frequency of that nucleotide to be in that position in real dataset and obtained a total score for each simulated read. Accordingly, best scoring 10 million reads were selected for both species from the simulated dataset with using `filter_syn.go` (Akkose et al. 2019)

### Downstream Analysis

To calculate nucleotide frequencies of excised oligomers we first converted aligned Xr-seq reads to `fasta` format using `bedtools getfasta`. Then using `frequency.go` (Akkose et al. 2019) we calculated nucleotide frequency of predominant oligomer (26nt) for each primate and damage type.

Command line used to get excised oligomer lengths: `awk ‘{print $3-$2}’ ${SAMPLE}_cutadapt_sorted.bed | sort -k1,1n | uniq -c | sed ‘s/\s\s*/ /g’ | awk ‘{print $2”\t”$1}’`

Statistical analyses of immunoslot blot repair assays and clonogenic survival assay have been performed with Welch Two Sample t-test by using `t-test.R` (Akkose et al. 2019).

For both genomes all annotated protein-coding genes were retrieved and genes that are closer to each other with less than 20kb were filtered out. We used 5277 and 3366 number of genes for human and mouse lemur, respectively. 10 kb upstream and downstream of TSS and TES were divided into 100 bp windows. XR-seq reads falling onto each bin were normalized to RPKM with `tcr.py` (Akkose et al. 2019) Then XR-seq RPKM values were normalized by the RPKM values derived by the mapped reads onto random genomic sites which are prepared using `bedtools shuffle` for protein-coding genes that are not closer to each other with less than 20kb. Statistical significance of TCR profiles was estimated using Mann-Whitney U-test (Mann and Whitney 1947). Statistical analyses were performed using `mann-w.py` (Akkose et al. 2019) with SciPy (Virtanen et al. 2020).

We used `regions_rpkm.go` (Akkose et al. 2019) to calculate RPKM values of XR-seq, RNA-seq, and simulated XR-seq for each overlapping region. Regions with no XR-seq or RNA-seq reads were filtered out.

Circos plot for comprehensive summarization of all findings has been generated with `circosPlot.R` (Akkose et al. 2019) by using R package `circlize`. (Gu et al. 2014)

PCA plot has been generated from read counts with `plotPCA.R` (Akkose et al. 2019), while applying variance stabilizing transformation as described in DESeq2 (Love et al. 2014) vignette.

Plots were prepared using `plotly` and `ggplot2` (Villanueva and Chen 2019). The manuscript was written using Manubot (Himmelstein et al. 2019).

## Data Access

All raw and processed sequencing data generated in this study have been submitted to the NCBI Gene Expression Omnibus (GEO; https://www.ncbi.nlm.nih.gov/geo/) under accession number GSE145883.

## Acknowledgments

This work is supported by National Institutes of Health grants GM118102 and ES027255 to AS. ADY was supported by NSF DEB-1354610, the John Simon Guggenheim Foundation, and the Alexander von Humboldt Foundation. OA is supported by EMBO Installation Grant (funded by TUBITAK: The Scientific and Technological Research Council of Turkey) and by TUBITAK 2232 International Fellowship for Outstanding Researchers Program. OA received a research grant as a recepient of Young Scientist Award (BAGEP) given by Science Acadamey, Turkey. OA also recieved personal research funds by Sabanci University. We would like to thank Dr. Beth A Sullivan for the construct to make the TERT-immortalized lines.

## Authors’ contributions

OA, AS, LL-B and ADY conceived of the project and designed the experiments. Mouse lemur tissue was acquired by ADY and PAL, cell lines were prepared by AB providing reagents for immortalization (hTERT construct). LL-B performed experimental assays, XR-seq and RNA-seq experiments. VOK, UA and ZK performed computational analyses. OA interpreted the data and wrote the first draft. All authors have read, revised and approved the final manuscript.

